# CK1δ-Dependent SNAPIN Dysregulation Drives Lysosomal Failure in HIV-1 Vpr–Exposed Neurons: A Targetable Mechanism in HAND

**DOI:** 10.1101/2025.07.11.664248

**Authors:** Bassel E. Sawaya, Maryline Santerre

## Abstract

HIV-associated neurocognitive disorders (HAND) persist in nearly 40% of virally suppressed individuals despite antiretroviral therapy (ART). Lysosomal dysfunction has emerged as a key contributor to HAND pathogenesis, yet the molecular mechanisms linking chronic HIV exposure to impaired neuronal degradation remain incompletely defined. Here, we identify HIV-1 Viral Protein R (Vpr) as a driver of lysosomal acidification failure, clustering, and degradative impairment in neurons. We uncovered casein kinase 1 delta (CK1δ) as a central mediator of this dysfunction, acting via phosphorylation of the adaptor protein SNAPIN. Vpr-induced CK1δ activation leads to hyperphosphorylation of SNAPIN, disrupting lysosomal positioning and motility. These defects are rescued by selective CK1δ inhibition, which restores lysosomal acidification, positioning, and mitophagy. Our findings define a novel Vpr–CK1δ–SNAPIN axis contributing to HAND and highlight lysosomal transport as a targetable mechanism in neurodegeneration.

**Graphical Abstract:** 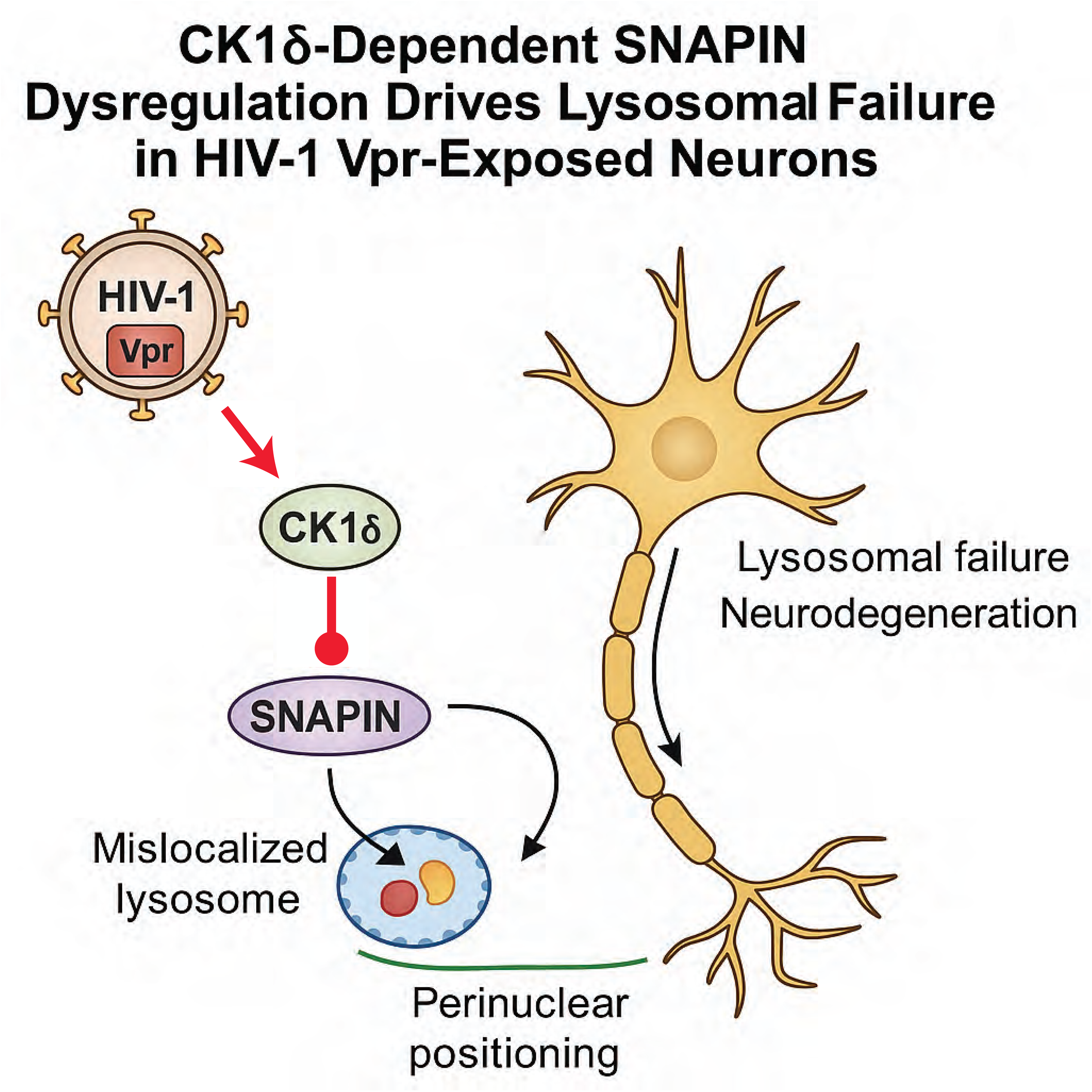

## INTRODUCTION

HIV-associated neurocognitive disorders (HAND) remain a significant clinical challenge affecting approximately 40-50% of People Living With HIV (PLWH) despite effective viral suppression with combination antiretroviral therapy (cART) ^1^. The spectrum of HAND encompasses conditions ranging from asymptomatic neurocognitive impairment (ANI) to mild neurocognitive disorder (MND) and the more severe HIV-associated dementia (HAD) ^2^. While the incidence of severe dementia has declined with modern antiretroviral therapies, milder forms of cognitive impairment persist and significantly impact quality of life ^3^. The persistence of these neurological complications despite undetectable viral loads in peripheral blood suggests complex pathogenic mechanisms beyond active viral replication ^4–6^.

Lysosomes play essential roles in neuronal homeostasis through the degradation of damaged organelles, protein aggregates, and other cellular waste products ^7^. Disruption of lysosomal acidification, positioning, or enzymatic function can lead to the accumulation of neurotoxic substrates, amplifying oxidative stress and inflammation. This phenotype is observed in Alzheimer’s disease, Parkinson’s disease^8, 9^, and increasingly in HAND. Autopsy studies of HAND patients reveal characteristic morphological alterations in endolysosomes and mitochondria, suggesting that these organelles are targets of HIV-induced neuronal injury ^10–13^.

HIV-1 Viral Protein R (Vpr) is a 96-amino acid accessory protein that continues to be detected in the cerebrospinal fluid of PLWH despite long-term antiretroviral therapy^14, 15^. Vpr exhibits remarkable functional versatility, affecting multiple cellular processes including cell cycle regulation, nuclear transport, and immune function ^16–18^. In the context of HIV neuropathogenesis, Vpr has been demonstrated to cause direct neurotoxicity by inducing mitochondrial dysfunction and oxidative stress ^19, 20^ Furthermore, extracellular Vpr decreases neuronal ATP levels ^21^ and disrupts intracellular redox balance, ultimately affecting neuronal health ^22–24^.

Recent investigations have uncovered a previously unrecognized role of Vpr in lysosomal biology. Specifically, Vpr has been shown to impair lysosomal clearance in neurons by decreasing lysosomal acidification and deregulating lysosomal positioning ^25^. These alterations result in the accumulation of substrates like alpha-synuclein (SNCA), potentially establishing mechanistic links between HIV infection and neurodegenerative processes observed in aging and Parkinson’s disease. However, the precise molecular mechanisms by which Vpr mediates these effects on lysosomal function remain incompletely defined.

Casein kinase 1 delta (CK1δ), a serine/threonine protein kinase, regulates diverse cellular processes including membrane trafficking, circadian rhythm, cell cycle progression, and autophagy ^26^. CK1δ dysregulation has been linked to Alzheimer’s disease (AD), where its aberrant activity alters tau phosphorylation, promotes neuroinflammation, and disrupts proteostasis ^27^.

Previous studies have identified SNAPIN as an interaction partner of CK1δ through yeast two-hybrid screening ^28^. SNAPIN serves as a critical adaptor protein that facilitates the recruitment of late endosomes to the dynein motor complex^29^, enabling retrograde transport along microtubules and subsequent lysosome maturation ^30^. This process is essential for maintaining proper lysosomal positioning and function in neurons, where polarized trafficking must occur over considerable distances ^31, 32, 33^. Dysregulation of this trafficking axis could therefore compromise neuronal clearance mechanisms in both infectious and non-infectious neurodegenerative contexts.

Phosphorylation events tightly regulate SNAPIN’s interaction with the dynein motor complex ^34^ and consequently affect lysosomal trafficking ^35^ Multiple kinases, including DYRK3 ^36^ and p38α- MAPK ^37^, have been shown to phosphorylate SNAPIN and modulate its functions. SNAPIN also interacts with components of the BORC/BLOC-1 complexes, multisubunit assemblies that regulate lysosomal positioning and the biogenesis of lysosome-related organelles ^38, 39^. Disruption of these interactions can lead to aberrant lysosomal distribution and compromised degradative capacity.

Mitophagy, the selective degradation of damaged mitochondria by autophagy, represents an essential quality control mechanism in neurons ^40^. HIV infection has been shown to alter mitochondrial dynamics in the CNS, with evidence suggesting that imbalances in mitochondrial fission/fusion contribute to neurodegeneration in HAND ^41^.

Given the established roles of both Vpr and the CK1δ-SNAPIN axis in lysosomal positioning and function, along with the known impact of HIV infection on mitochondrial quality control processes, we sought to investigate the potential relationship between these pathways in the context of HIV-associated neurodegeneration. Specifically, we hypothesized that HIV-1 Vpr might disrupt lysosomal positioning and degradative capacity through activation of CK1δ, leading to aberrant phosphorylation of SNAPIN. Our investigations into this Vpr–CK1δ–SNAPIN signaling axis using complementary molecular, imaging, and proteomic approaches provide novel insights into the mechanisms by which chronic HIV exposure contributes to neuronal damage and identify CK1δ as a potential therapeutic target for HAND.

## RESULTS

### 1. HIV-1 Vpr Drives Lysosomal Overload in Neurons via Impaired Degradation of Diverse Cargoes

To determine whether HIV-1 Vpr disrupts lysosomal degradation in neurons, we treated SH-SY5Y cells differentiated into neurons with recombinant Vpr (8 nM) and assessed the clearance of diverse lysosomal cargoes by confocal microscopy. We observed a marked accumulation of lipid droplets, as revealed by BODIPY staining, which colocalized with the lysosomal marker LAMP1 (**Fig. 1A**). Similarly, mitochondrial remnants retained MitoTracker signal even after CCCP-induced depolarization, indicating impaired mitophagy. These undegraded mitochondria accumulated in LAMP1⁺ compartments following Vpr exposure (**Fig. 1B**).

**Figure 1.**
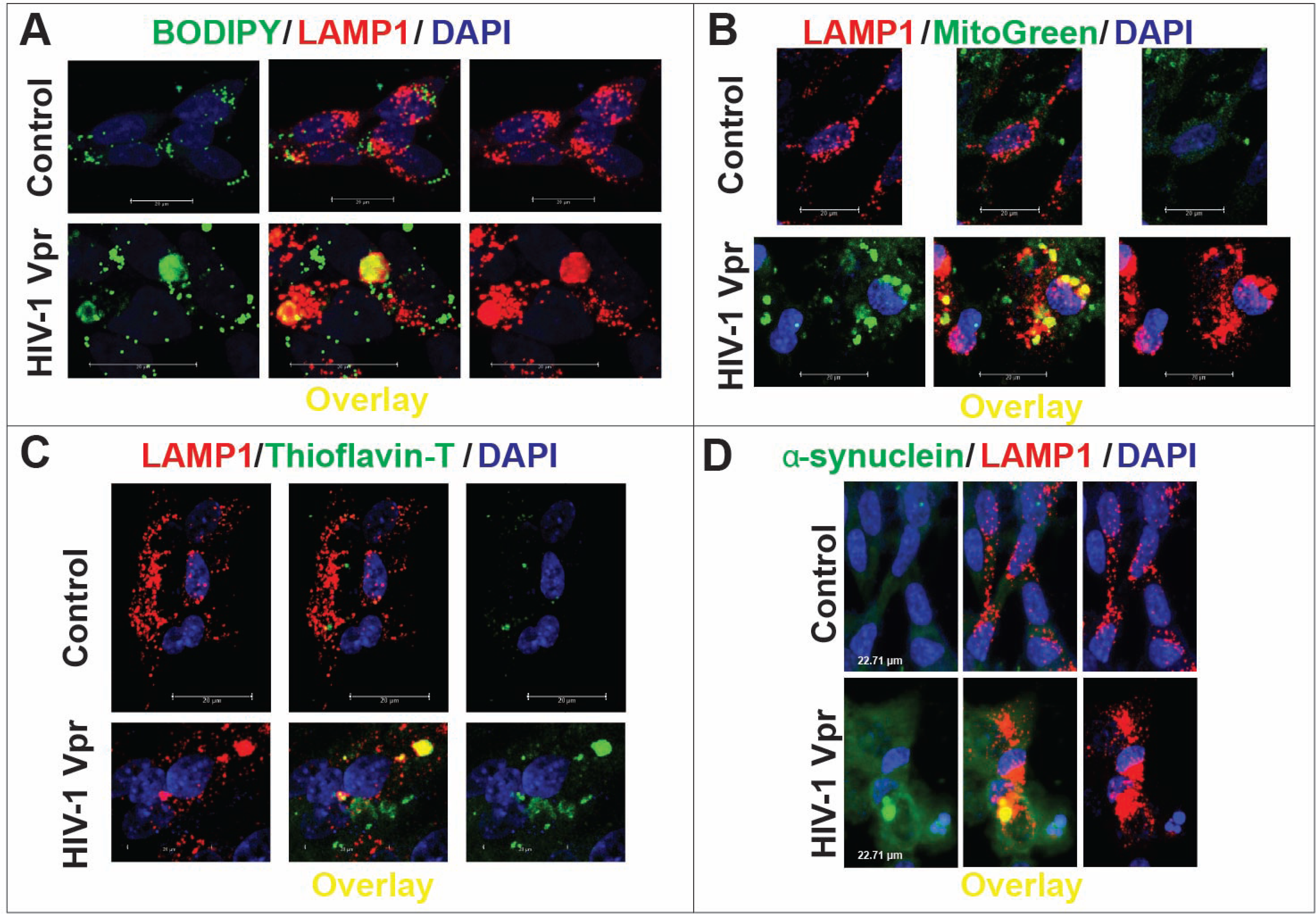
HIV-1 Vpr impairs lysosomal degradation, leading to the accumulation of diverse cargoes in neuronal lysosomes. SH-SY5Y cells stably expressing LAMP1-mCherry differentiated into neurons were treated with recombinant HIV-1 Vpr (8 nM) and analyzed by confocal microscopy. **(A)** Lipid accumulation was detected by BODIPY staining (green), which showed increased colocalization with the lysosomal marker LAMP1 (red) in Vpr-treated cells. **(B)** Mitochondrial remnants labeled with MitoTracker (green) persisted in lysosomes following Vpr exposure, indicating defective mitophagy. **(C)** Thioflavin-T staining (green) revealed increased amyloid-like deposits within LAMP1⁺ lysosomes in Vpr-treated neurons. **(D)** α-synuclein (green) also showed enhanced lysosomal association upon Vpr exposure. DAPI (blue) labels nuclei. Across all cargo types, Vpr increased lysosomal entrapment, as confirmed by quantitative colocalization analysis (Manders’ coefficients), indicating impaired degradation. Scale bars: 20 µm.

In addition to metabolic substrates, we examined the fate of aggregated proteins. Thioflavin-T staining revealed increased amyloid-like deposits in Vpr-treated neurons, again colocalizing with lysosomes (**Fig. 1C**). Moreover, α-synuclein (SNCA), a protein implicated in neurodegeneration, showed enhanced lysosomal association following Vpr exposure (**Fig. 1D**). Across all conditions, quantitative image analysis confirmed significantly elevated Manders’ colocalization coefficients between each cargo and LAMP1, consistent with their impaired degradation and entrapment within lysosomes.

Together, these data demonstrate that HIV-1 Vpr impairs lysosomal degradative capacity, leading to the accumulation of undigested lipids, mitochondria, and protein aggregates within neuronal lysosomes—a process that may contribute to neurodegeneration in HAND.

### 2. HIV-1 Induces SNAPIN Mislocalization Across Neuronal Populations, with Marked Aggregation in Cerebellar and Purkinje Neurons

To investigate whether HIV-1 Vpr affects the subcellular distribution of SNAPIN in vivo, we performed immunohistochemical analysis of brain sections from control and HIV-1 transgenic rats (**Fig**. **2**). In control F344/N rats, SNAPIN was diffusely expressed in the cytoplasm of Purkinje cells and molecular layer interneurons, in the cerebellar cortex, without evidence of clustering. In contrast, HIV-1 transgenic brains displayed markedly increased SNAPIN signal, characterized by perinuclear clustering and punctate aggregation within Purkinje neurons and adjacent interneurons. Similar SNAPIN accumulation was also observed in extracerebellar regions, including the locus coeruleus, suggesting widespread dysregulation (data not shown). These findings indicate that HIV-1 Vpr drives abnormal SNAPIN accumulation and mislocalization in neurons, particularly in vulnerable cerebellar cell types.

**Figure 2.**
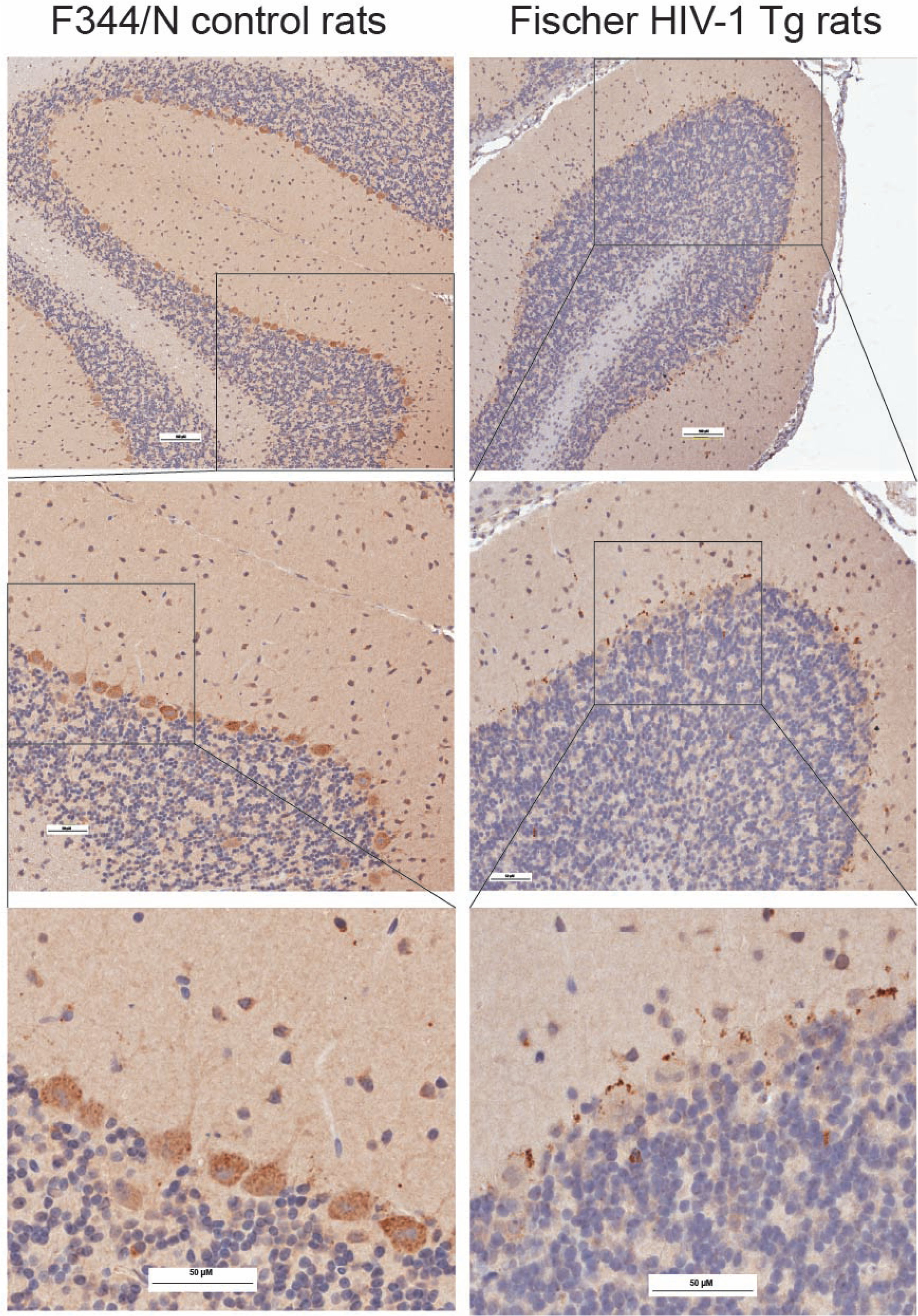
HIV-1 induces SNAPIN mislocalization and aggregation in transgenic rats neurons. Immunohistochemical analysis of brain sections (here the cerebellum is shown) from control F344/N rats (left) and HIV-1 transgenic Fischer rats (right) revealed altered SNAPIN distribution in vivo. In control animals, SNAPIN was diffusely localized in the cytoplasm of Purkinje cells and molecular layer interneurons. In contrast, HIV-1 Tg rats exhibited increased SNAPIN accumulation, with pronounced perinuclear clustering and punctate aggregates, particularly within Purkinje neurons. Top panels show low-magnification views of the cerebellar cortex. Middle panels present higher magnification of the boxed regions in the top panels. Bottom panels show further magnified views of the boxed areas from the middle panels, highlighting perinuclear SNAPIN clustering in Purkinje neurons. Scale bars: 100 and 50 µm.

### 3. HIV-1 Vpr Disrupts SNAPIN Serine 50 Phosphorylation, Impairing Lysosomal Trafficking in Neurons

We previously reported that Vpr reduces SNAPIN expression, contributing to lysosomal mislocalization and impaired degradation in neurons ^25^. Here, we investigated whether HIV-1 Vpr also alters SNAPIN through posttranslational modifications. Immunoprecipitation followed by immunoblotting revealed that Vpr increases SNAPIN phosphorylation on serine residues while decreasing its O-GlcNAcylation—a modification associated with protein stability and trafficking function (**Fig. 3A**).

**Figure 3.**
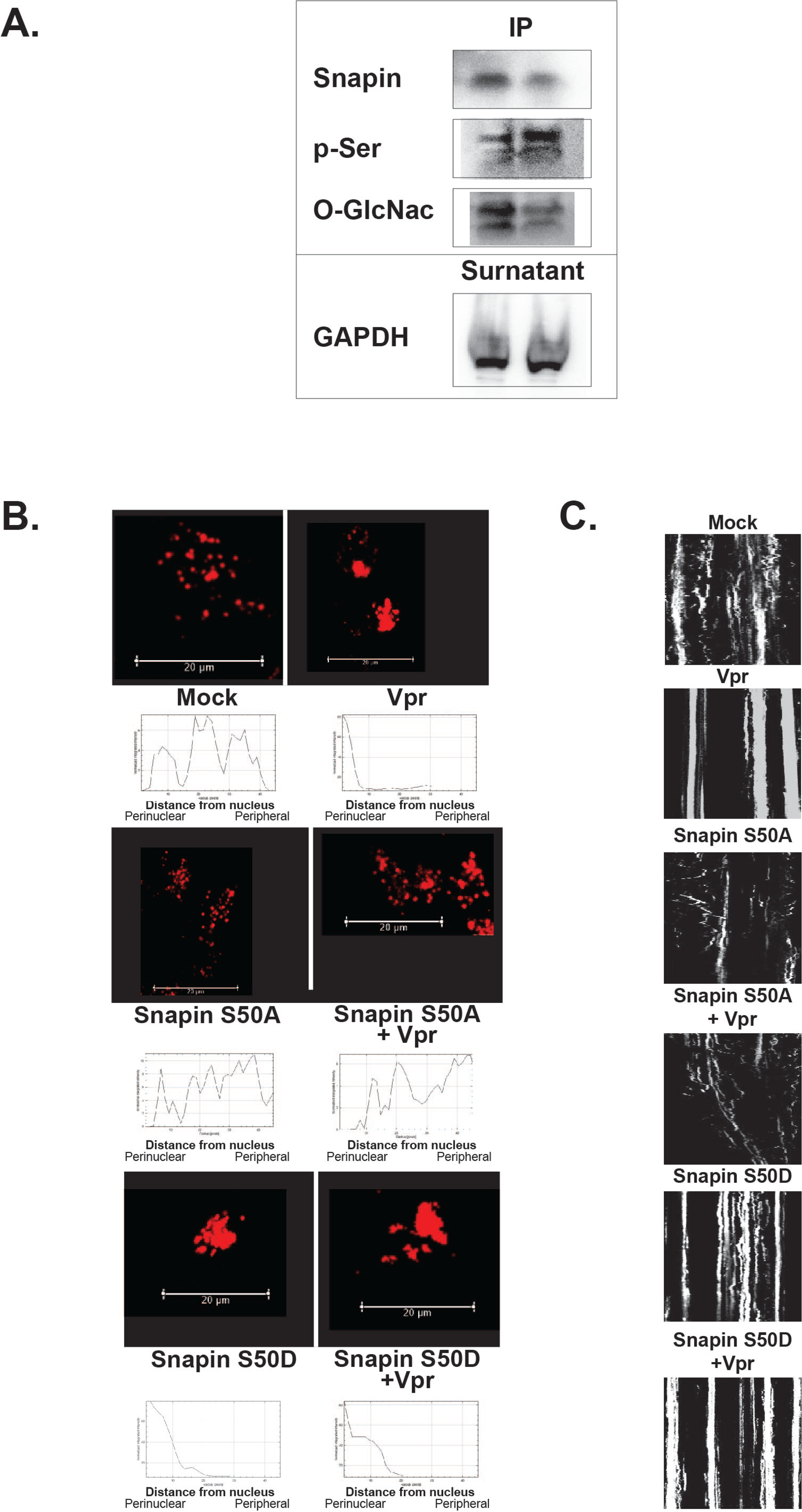
HIV-1 Vpr alters SNAPIN posttranslational modifications and disrupts lysosomal trafficking via serine 50. (A) Immunoprecipitation and Western blot of SNAPIN in Vpr-treated neurons revealed increased serine phosphorylation (p-Ser) and decreased O-GlcNAcylation, indicating posttranslational modulation. GAPDH in the supernatant served as a loading control. (B) Confocal images and radial intensity plots show lysosomal clustering in neurons expressing SNAPIN WT and mutants with or without Vpr. Vpr induced perinuclear lysosome accumulation that was mimicked by phosphomimetic S50D and prevented by the non-phosphorylatable S50A mutant. Scale bars: 20 µm (C) Kymograph analysis of LAMP1⁺ vesicle movement demonstrates that Vpr and S50D both impair lysosomal motility, whereas S50A preserves trafficking, and is insensitive to Vpr effect.

To assess the functional relevance of SNAPIN phosphorylation, we systematically mutated all known phosphorylation sites identified in PhosphoSitePlus (www.phosphosite.org) and examined lysosome positioning in neurons. Of all the mutants tested, only alteration of serine 50 produced a distinct lysosomal phenotype, prompting focused analysis of SNAPIN serine 50 phospho-mutants to further characterize its role in lysosomal regulation. Vpr exposure induced prominent perinuclear clustering of lysosomes with reduced overlap with peripheral LAMP1⁺ vesicles, consistent with our previous findings ^25^, and this phenotype was recapitulated by the phosphomimetic S50D mutant (**Fig. 3B**). In contrast, the non-phosphorylatable S50A mutant preserved normal lysosomal distribution even in the presence of Vpr. Kymograph analysis in SH-SY5Y cells confirmed that lysosomal motility was impaired in both Vpr-exposed and S50D-expressing neurons, whereas S50A maintained directional trafficking (**Fig. 3C**). Together, these results identify serine 50 as a critical regulatory site through which Vpr impairs SNAPIN function, leading to defective lysosomal positioning and motility in neurons.

### 4. HIV-1 Vpr Induces CK1δ Expression, Leading to SNAPIN Phosphorylation

Given that SNAPIN phosphorylation at serine 50 disrupts lysosomal positioning and motility in Vpr-exposed neurons, we next sought to identify the upstream kinase responsible. Using a pharmacological screen, we tested whether kinase inhibition could rescue Vpr-induced trafficking defects. Among candidates, only the CK1δ inhibitor LH846 restored lysosomal function, prompting us to focus on casein kinase 1 delta (CK1δ)—a serine/threonine kinase known to regulate vesicle transport and neuronal signaling.

#### HIV-1 Vpr Upregulates CK1δ Expression

To explore whether CK1δ is regulated by Vpr, we assessed CK1δ expression by immunofluorescence (**Fig. 4**). Vpr exposure resulted in increased CK1δ staining, which was partially reversed by LH846 treatment. Quantification confirmed that Vpr upregulates CK1δ expression in neurons, suggesting that Vpr may directly or indirectly activate this kinase as part of its pathogenic program.

**Figure 4.**
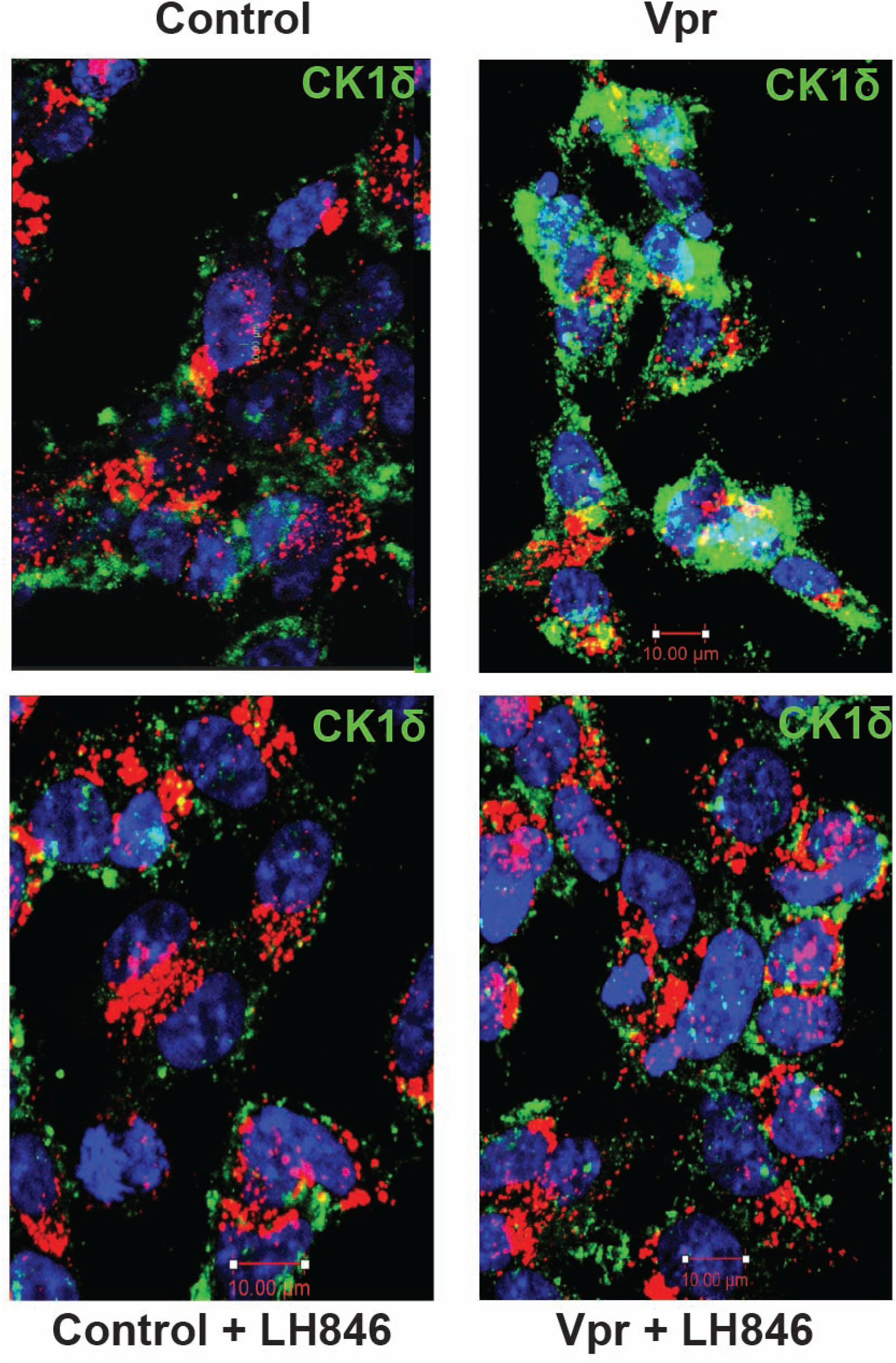
HIV-1 Vpr upregulates CK1δ expression in neurons. (A) Immunofluorescence staining of CK1δ (green) in SH-SY5Y-derived neurons treated with vehicle (Control), Vpr, LH846, or Vpr + LH846 or DMSO. CK1δ signal intensity increased following Vpr exposure and was partially reduced by LH846 treatment. Nuclei are labeled with DAPI (blue); LAMP1 (red) marks lysosomes. Scale bars: 10 µm (B) Quantification of CK1δ fluorescence intensity per cell confirms a significant increase in Vpr-treated neurons, with partial normalization by CK1δ inhibition (mean ± SEM, n = 100 cells minimum per condition).

#### CK1δ Inhibition Restores Lysosomal Positioning

To determine whether CK1δ contributes to lysosomal mispositioning, we assessed the spatial distribution of LAMP1⁺ vesicles in neurons treated with Vpr, with or without the CK1δ inhibitor LH846 (**Fig. 5**). Control neurons displayed broadly distributed lysosomes, whereas Vpr exposure induced perinuclear clustering. LH846 restored peripheral lysosomal distribution, while co-expression of phosphomimetic SNAPIN (S50D) blocked this rescue. This finding underscores the functional importance of serine 50 in mediating CK1δ-dependent lysosomal clustering, linking CK1δ activity directly to SNAPIN phosphorylation at this site.

**Figure 5.**
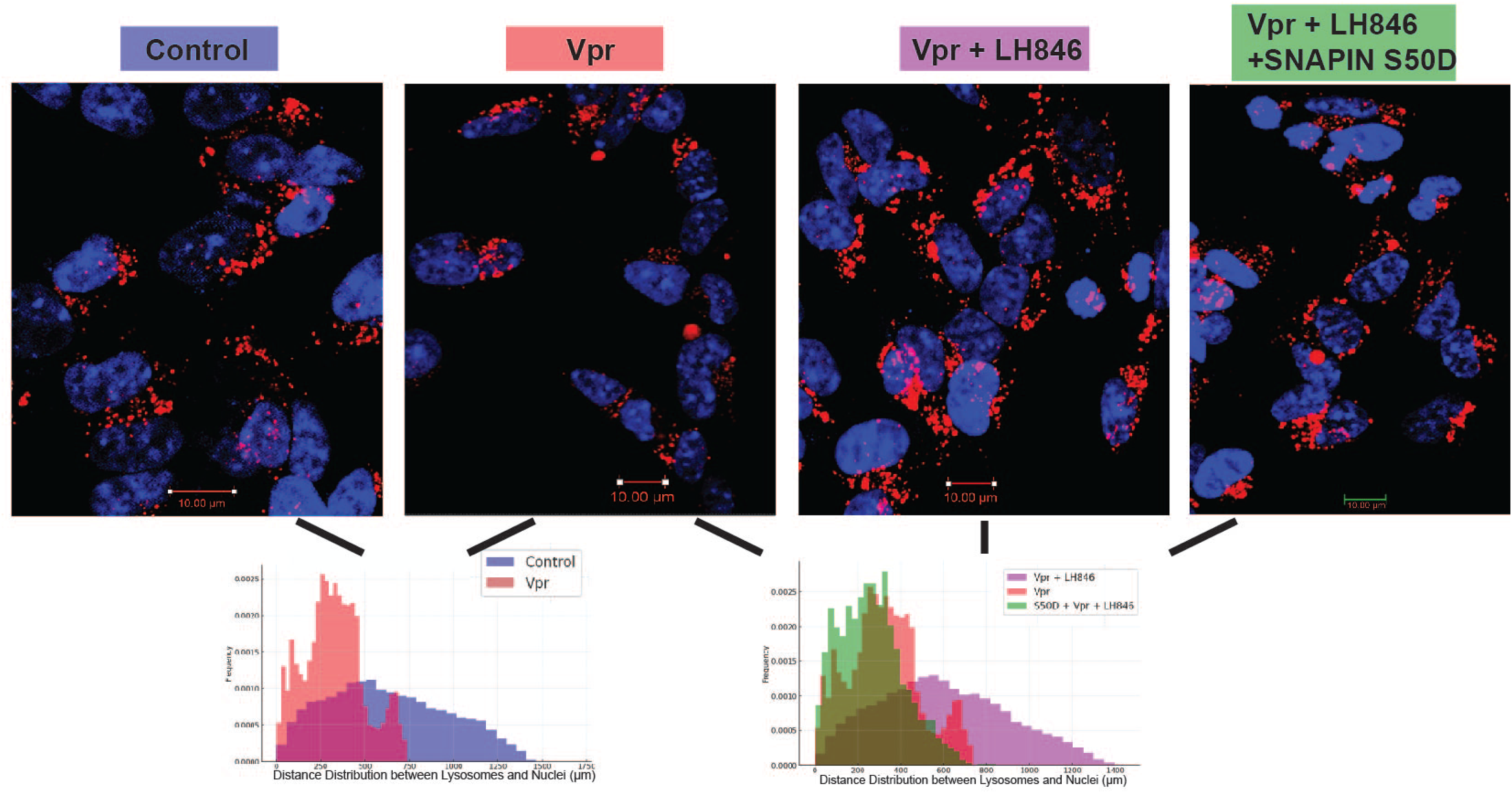
CK1δ inhibition restores lysosomal distribution disrupted by HIV-1 Vpr. SH-SY5Y cells stably expressing LAMP1-mCherry were transfected with SNAPIN WT or phosphomimetic SNAPIN S50D, differentiated into neurons for 4 days and treated 24h with Vpr ± CK1δ inhibitor LH846 (1.5μM). LAMP1⁺ lysosomes (red) and nuclei (blue, DAPI) were imaged by confocal microscopy. Lysosomal spatial distribution was quantified using radial intensity profiles from nuclear centers in ImageJ (ImageJ Radial Profile plugin), and data were presented as normalized histograms of different colors to facilitate visual comparison between groups.

#### CK1δ Inhibition Restores SNAPIN Localization

We next tested whether CK1δ inhibition could reverse Vpr-induced SNAPIN mislocalization (**Fig. 6A**). In Vpr-treated neurons, SNAPIN accumulated in clustered perinuclear aggregates and showed increased colocalization with LAMP1⁺ lysosomes—potentially explaining the overall reduction in detectable SNAPIN levels. Treatment with the CK1δ inhibitor LH846 restored a more diffuse cytoplasmic distribution of SNAPIN and significantly reduced its lysosomal association, as quantified by Manders’ coefficient (**Fig. 6B**). These results indicate that CK1δ activity mediates Vpr-driven SNAPIN mislocalization and promotes its aberrant accumulation within lysosomes.

**Figure 6.**
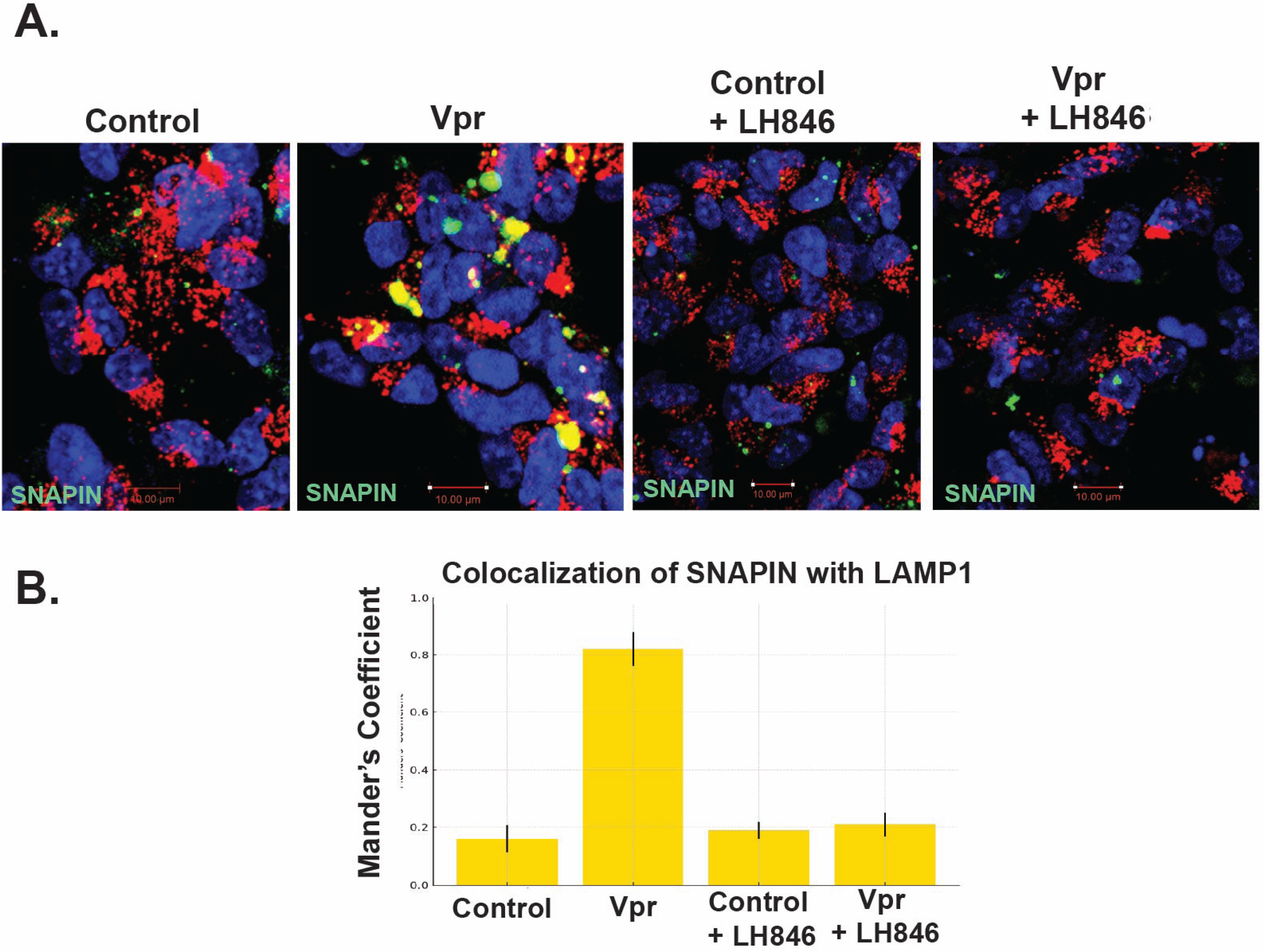
CK1δ inhibition restores SNAPIN distribution and rescues lysosomal positioning. SH-SY5Y cells stably expressing LAMP1-mCherry were transfected with SNAPIN-GFP, differentiated into neurons for 4 days and treated 24h with Vpr ± CK1δ inhibitor LH846 (1.5μM). LAMP1⁺ lysosomes (red), SNAPIN-GFP (Green) and nuclei (blue, DAPI) were imaged by confocal microscopy. Quantification (bottom panel) of Manders’ colocalization coefficient between SNAPIN and LAMP1 confirms a significant increase with Vpr that is reversed by CK1δ inhibition (mean ± SEM; n = 100 cells minimum). Scale bar: 10 μm.

### 5. CK1δ Inhibition Restores Lysosomal Acidification and Mitophagy in Vpr-Exposed Neurons

To determine whether CK1δ inhibition rescues lysosomal function downstream of Vpr, we first assessed lysosomal pH using a dual-emission pH-sensitive LC3 reporter (pHluorin-mCherry-LC3) (**Fig. 7**). Vpr exposure significantly increased the green/red (G/R) fluorescence ratio, consistent with lysosomal deacidification (**Fig. 7A**). Treatment with the CK1δ inhibitor LH846 restored acidification to near-baseline levels, indicating that CK1δ activity contributes to Vpr-induced lysosomal alkalinization.

**Figure 7.**
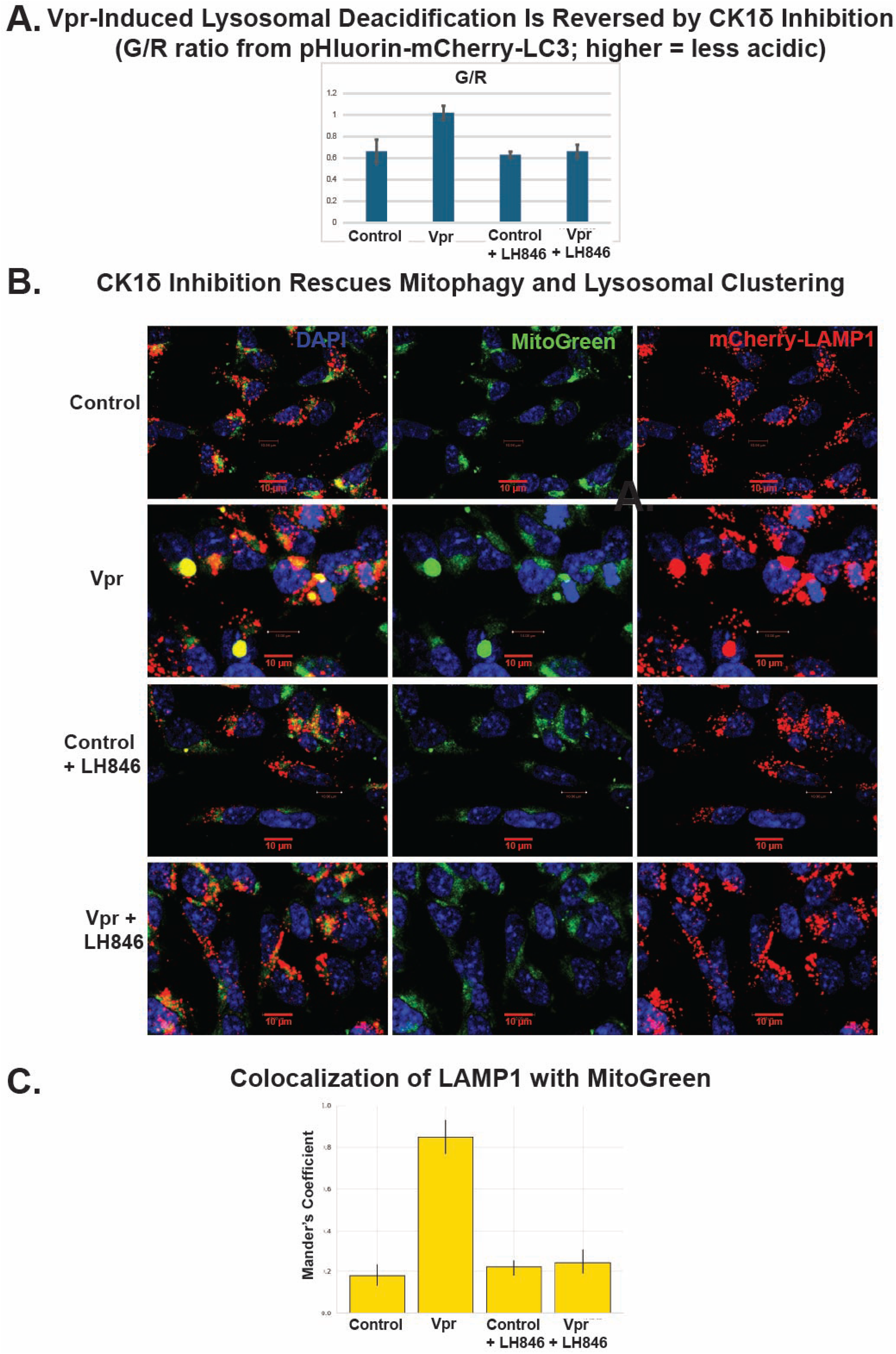
CK1δ inhibition restores lysosomal acidification and mitophagy in Vpr-exposed neurons. (A) Quantification of lysosomal pH using the pH-sensitive dual-emission LC3 reporter (pHluorin-mCherry-LC3). Vpr treatment significantly increased the green/red (G/R) fluorescence ratio, indicating lysosomal deacidification, which was reversed by the CK1δ inhibitor LH846 (mean ± SEM; n = 100 cells minimum). (B) Confocal images of SH-SY5Y-derived neurons stably expressing mCherry-LAMP1 and stained with MitoBright LT Green (Abbreviated as MitoGreen) (Dojindo) show that Vpr induces accumulation of mitochondria colocalizing with lysosomes, consistent with impaired mitophagy. LH846 treatment reduced mitochondrial buildup and restored spatial segregation between mitochondria and lysosomes. (C) Quantification of LAMP1–MitoGreen colocalization using Manders’ coefficient confirms elevated overlap in Vpr-exposed neurons and significant rescue by LH846. Scale bars: 10 μm.

We next evaluated mitophagy and mitochondrial degradation by co-expressing a mitochondrial marker (MitoBright LT Green, simplified as MitoGreen) and a lysosomal marker (mCherry- LAMP1) (**Fig. 7B**). In control neurons, damaged mitochondria were efficiently cleared, with minimal overlap between MitoGreen and LAMP1. In contrast, Vpr-exposed neurons displayed large accumulations of undegraded mitochondria that colocalized with LAMP1⁺ compartments, consistent with impaired mitophagy and lysosomal degradation. LH846 treatment markedly reduced MitoGreen retention and decreased LAMP1–mitochondria colocalization, as quantified by Manders’ coefficient (**Fig. 7C**).

Together, these results demonstrate that CK1δ inhibition restores lysosomal acidification and degradative capacity in Vpr-exposed neurons, reversing mitochondrial and autophagic defects associated with SNAPIN misregulation.

## DISCUSSION

Despite viral suppression with modern antiretroviral therapy, HIV-associated neurocognitive disorders (HAND) persist in a substantial subset of people living with HIV ^1^, underscoring the need to uncover virus-mediated, non-replicative mechanisms of neuronal dysfunction. Our study identifies a previously unrecognized signaling axis—Vpr–CK1δ–SNAPIN—that links HIV-1 exposure to lysosomal dysfunction, providing mechanistic insight into how chronic viral protein activity can disrupt organellar homeostasis and drive neurodegeneration.

We demonstrate that exposure to HIV-1 Vpr impairs lysosomal degradation by inducing the accumulation of undigested cargoes—including lipids, mitochondria, amyloid aggregates, and α- synuclein—within LAMP1⁺ compartments. These findings align with reports of endolysosomal abnormalities in HAND brain tissue^10–13^ and extend them by identifying SNAPIN misregulation as a central mediator of lysosomal trafficking failure. Specifically, we show that Vpr increases CK1δ expression in neurons, leading to SNAPIN hyperphosphorylation at serine 50, and mislocalization, possibly leading to functional uncoupling from motor complexes ^29^. This disruption in SNAPIN function impairs lysosomal positioning, motility, and degradative capacity—hallmarks of lysosomal stress previously reported in both HIV- and aging-associated neurodegenerative contexts^10^.

Importantly, our phosphomimetic SNAPIN mutant (S50D) recapitulates the Vpr-induced defects in lysosomal positioning and motility, while the non-phosphorylatable S50A mutant is protective. Pharmacological inhibition of CK1δ with LH846 reverses these effects—restoring SNAPIN localization, lysosomal acidification, and mitophagy—highlighting CK1δ as both an effector of Vpr toxicity and a promising therapeutic target in HAND. We propose a working model in which Vpr acts upstream of CK1δ, initiating a cascade that impairs neuronal clearance mechanisms through SNAPIN phosphorylation and lysosomal immobilization.

Our in vivo immunohistochemistry data extend these findings to the cerebellum, revealing that HIV-1 transgenic rats exhibit SNAPIN aggregation specifically in Purkinje cells and interneurons^42^—cell types increasingly recognized for their role in fine motor control^43^, autonomic regulation^44^, and sleep architecture^45^. Purkinje neurons, in particular, are highly polarized and metabolically demanding, rendering them especially vulnerable to defects in organelle trafficking. Lysosomal dysfunction in these cells could therefore contribute to both the motor abnormalities (e.g., tremor, ataxia) and sleep disturbances frequently observed in HAND patients. Notably, neuroimaging studies in HIV+ individuals have reported significant reductions in frontal, brainstem, and cerebellar volumes in patients on integrase strand transfer inhibitor (INSTI)-based therapy compared to non-INSTI users^46^, supporting a broader relevance of cerebellar vulnerability in HIV-associated neurodegeneration. Thus, our identification of a lysosome-targeting mechanism active in Purkinje neurons provides a potential cellular basis for diverse clinical symptoms in HAND.

Mechanistically, our work builds on prior evidence linking CK1 family kinases to vesicle trafficking, proteostasis, and circadian biology^47^. However, we are the first to implicate CK1δ as a Vpr-responsive kinase that phosphorylates SNAPIN and impairs lysosomal trafficking in neurons. This adds to a growing recognition that viral proteins can hijack host kinases to induce long-term cellular dysfunction, even in the absence of active viral replication^48, 49^. It also raises the possibility that CK1δ functions as a stress amplifier, integrating Vpr toxicity with broader disruptions in organellar homeostasis and clearance.

In conclusion, we define a novel Vpr–CK1δ–SNAPIN signaling axis that underlies lysosomal dysfunction in neurons and contributes to HAND pathogenesis. These findings position CK1δ as a promising therapeutic target to restore neuronal degradative function in the context of chronic HIV exposure. Future studies will be needed to determine whether CK1δ inhibition can reverse cognitive deficits in vivo and to explore whether this axis plays a broader role in other neurodegenerative or lysosomal storage disorders where SNAPIN function is compromised.

## Limitations of the Study

While this study establishes a mechanistic link between HIV-1 Vpr, CK1δ activation, and SNAPIN-mediated lysosomal dysfunction, several limitations should be considered. First, our use of neuronal cell lines limits interpretation of circuit-level or behavioral outcomes; future studies employing in vivo models with cognitive or motor assessments will be necessary to define functional relevance. Second, while LH846 effectively rescued Vpr-induced defects in vitro, its pharmacokinetic properties, brain penetrance, and long-term safety profile remain uncharacterized. Thus, further evaluation of CK1δ inhibition in animal models will be essential to determine translational feasibility.

## Acknowledgments

This work was supported by an NIH-NIA grant AG054411 awarded to BES, and an NIH-NIA grant R56AG082530 to MS. We thank Kathy Q. Cai (Fox Chase Cancer Center) for performing the immunohistochemistry and Dr. Rosemarie M. Booze (University of South Carolina) for generously providing the rat brain tissue. The authors declare use of generative AI tools: During the preparation of this work, the authors used ChatGPT to improve clarity, flow, and readability. AI was used solely for language editing; all content was reviewed and verified by the authors, who take full responsibility for the final manuscript.

## MATERIALS AND METHODS

### Plasmids

pLAMP1-mCherry was a gift from Amy Palmer (Addgene plasmid #45147; http://n2t.net/addgene:45147; RRID:Addgene_45147)¹. FUGW-PK-hLC3 was a gift from Isei Tanida (Addgene plasmid #61460; http://n2t.net/addgene:61460; RRID:Addgene_61460)². The SNAPIN plasmid was a gift from Jing Pu, PhD³. pcDNA Vpr expression plasmid containing the CMV promoter expressing Vpr was previously described{Mahalingam, 1995 #819}.

### Cell Culture and Treatments

Human SH-SY5Y cells were differentiated into neurons using retinoic acid (10 µM) for 5 days prior to experiments {Kovalevich, 2021 #1529}. Cells were treated with recombinant HIV-1 Vpr protein (8 nM) for 24 hours. For kinase inhibition, LH846 (CK1δ inhibitor, Cayman Chemicals, 17686) was applied at 1.5 µM for the final 16 hours of treatment.

### Mutagenesis and Transfections

Site-directed mutagenesis with Q5® High-Fidelity DNA Polymerase (NEB, M0491S) was used to generate phosphomimetic (S50D) and non-phosphorylatable (S50A) SNAPIN mutants.

S50D: Forward: 5’ ACACGCCGTCAGAGAGGACCAGGTAGAGCT 3’, Reverse: 5’

AGCTCTACCTGGTCCTCTCTGACGGCGTGT 3’.

S50A: Forward: 5’ GTACACGCCGTCAGAGAGGCACAGGTAGAGCT 3’, Reverse: 5’

AGCTCTACCTGTGCCTCTCTGACGGCGTGTAC 3’.

### HA-Tagged Protein Immunoprecipitation

SH-SY5Y cells were co-transfected with SNAPIN-HA using Lipofectamine 3000 (Invitrogen, L3000015) and pcDNA-Vpr (200 ng) or empty vector and differentiated for 4 days{Kovalevich, 2021 #1529}. Cells were lysed in NP-40 lysis buffer (50 mM Tris-HCl pH 7.4, 150 mM NaCl, 1% NP-40, 1 mM EDTA) with protease and phosphatase inhibitors. Lysates were rotated at 4°C for 30 minutes, centrifuged (12,000 × g, 10 min), and incubated with 25 µL Pierce Anti-HA magnetic beads (88837) in TBS-T at room temperature for 30 min. Beads were washed and eluted in 2× Laemmli buffer (50°C, 10 min) for SDS-PAGE and immunoblotting. The immunoprecipitation is followed by immunoblotting with anti–phospho-serine (Santa Cruz, sc-81514) and O-GlcNAc antibodies (Santa Cruz, sc-74546). Bands were visualized with ECL.

### Immunocytochemistry and Imaging

SH-SY5Y cells stably expressing LAMP1-mCherry were differentiated into neurons and treated with recombinant HIV-1 Vpr (8 nM) for 24 hours and Mock (DMSO) or LH846 (1.5 µM), then fixed in 4% paraformaldehyde. **Lipid droplets** were visualized using BODIPY 493/503 (Thermo Fisher, D3922; 1 µg/mL, 15 min at room temperature). **Mitochondria** were labeled with MitoBright LT Green (Dojindo, MT10-12; 0.1 μmol/L, 30 min), with or without 10 µM CCCP (Sigma, C2759) for 3 hours to induce mitophagy. **Amyloid aggregates** were stained with Thioflavin T (Sigma, T3516; 1 µM, 15 min), and **α-synuclein** was detected by immunostaining with anti-SNCA antibody (Santa Cruz, sc-12767) followed by fluorescent secondary antibody. Nuclei were counterstained with DAPI (Sigma, D9542). CK1δ was detected using the Casein Kinase Iδ antibody (C-8; sc-55553, Santa Cruz Biotechnology) and a fluorescent secondary antibody. Coverslips were mounted with VECTASHIELD® Antifade Mounting Medium (Vector Laboratories, H-1000-10) and imaged using a Leica DM4000 confocal microscope. Colocalization with LAMP1 was quantified using Manders’ coefficient in ImageJ with the JACoP plugin.

**CK1δ:** Cells were fixed in 4% PFA and stained with antibodies against CK1δ (Proteintech, 14388- 1-AP). Imaging was performed using a Leica DM4000 confocal microscope. Colocalization was quantified using Manders’ coefficient in ImageJ.

**SNAPIN Localization:** SH-SY5Y neurons stably expressing LAMP1-mCherry were transfected with SNAPIN-GFP and treated with Vpr (8 nM, ±1.5 µM LH846 or DMSO, 24 h), then fixed in 4% PFA. Imaging used identical settings across conditions (Leica DM4000). ImageJ (Fiji) was used to project z-stacks and calculate Manders’ colocalization coefficient (JACoP plugin). ≥100 cells/condition across 3 replicates were analyzed.

**Mitophagy Assays**: Mitophagy was evaluated by imaging of neurons co-expressing MitoBright LT Green (Abbreviated as MitoGreen) (Dojindo) and LAMP1-mCherry (Confocal Leica DM4000), and quantified by colocalization analysis.

### Kymograph Analysis for Motility, and Lysosome Positioning Measurement

Live-cell imaging was performed on SH-SY5Y cells stably expressing LAMP1-mCherry to assess lysosomal motility and spatial distribution with a confocal microscope Leica DM4000 with constant 5% CO2 at 37°C degrees. Kymographs were generated using the Multi-Kymograph plugin in ImageJ (https://imagej.net/plugins/multi-kymograph) to quantify lysosome velocity and directionality. Lysosomal positioning relative to the nucleus was analyzed using the Radial Profile plugin in ImageJ (https://imagej.nih.gov/ij/plugins/radial-profile.html), which generated intensity plots from the nuclear center to the cell periphery.

### Lysosomal pH

Lysosomal acidification was assessed using a dual-emission pH-sensitive reporter construct (pHluorin-mCherry-LC3) expressed in SH-SY5Y neurons. Cell fluorescence is quantified by flow cytometry (Guava5HT, Millipore). Green is measured in the FL1 channel (Excitation wavelength 488 nm; Emission 515–545 nm), Red is measured in the FL2 channel (yellow). The median fluorescence value is determined for at least 10000 cells.

### IHC

Formalin-fixed, paraffin-embedded rat brain slides were processed at the Histopathology Facility (Fox Chase Cancer Center). Sections were incubated with antibodies and developed with DAB (Acros Organics, AC112090010) and counterstained with hematoxylin (Vector, H-3401). SNAPIN antibodies: Proteintech (10055-1-AP).

### Animal Studies

Brain slices were a gift from Dr. Rosemary Booze (University of South Carolina). All animal work was approved by the IACUC (Federal Assurance #D16-00028) and conducted in AAALAC-accredited facilities. 12-month-old Fischer HIV-1 Tg rats and F344/N controls were housed under a 12:12 light/dark cycle with ad libitum access to water and chow (#3000, Pro-Lab). Control rats were from Envigo (Indianapolis, IN); HIV-1 Tg rats were bred on-site.

